# High affinity interactions of Pb^2+^ with Synaptotagmin I

**DOI:** 10.1101/348748

**Authors:** Sachin Katti, Bin Her, Atul K. Srivastava, Alexander B. Taylor, Steve W. Lockless, Tatyana I. Igumenova

## Abstract

Lead (Pb) is a potent neurotoxin that disrupts synaptic neurotransmission. We report that Synaptotagmin I (SytI), a key regulator of Ca^2+^-evoked neurotransmitter release, has two high-affinity Pb^2+^ binding sites that belong to its cytosolic C2A and C2B domains. The crystal structures of Pb^2+^-complexed C2 domains revealed that protein-bound Pb^2+^ ions have holodirected coordination geometries and all-oxygen coordination spheres. The on-rate constants of Pb^2+^ binding to the C2 domains of SytI are comparable to those of Ca^2+^ and are diffusion-limited. In contrast, the off-rate constants are at least two orders of magnitude smaller, indicating that Pb^2+^ can serve as both thermodynamic and kinetic trap for the C2 domains. We demonstrate, using NMR spectroscopy, that population of these sites by Pb^2+^ ions inhibits further Ca^2+^ binding despite the existing coordination vacancies. Our work offers a unique insight into the bioinorganic chemistry of Pb(II) and suggests a mechanism by which low concentrations of Pb^2+^ ions can interfere with the Ca^2+^-dependent function of SytI in the cell.

## INTRODUCTION

Lead poisoning remains a pervasive public health problem, as illustrated by the recent outbreaks in the U.S. (Flint, Michigan) and abroad.^1, 2^ Lead exposure is especially detrimental in young children, resulting in serious neurodevelopmental and psychological disorders.^3–5^ The potency of Pb^2+^ ([Xe]-4*f*^14^5*d*^10^6*s*^2^) stems from its ability to cross the blood-brain barrier^6^ and preferentially target Zn^2+^ and Ca^2+^ coordination sites of biological macromolecules.^7–9^ The ability of Pb^2+^ to mimic these essential divalent metal ions results in disruption of cellular signaling, ion transport, and calcium homeostasis.^10–13^

The molecular mechanisms of Pb^2+^ neurotoxicity are not well understood. Several neuronal proteins associated with Ca^2+^ signaling have been implicated in Pb^2+^ toxicity (reviewed in^8^). Among them are the voltage-gated Ca^2+^ channels, where the putative mechanism is the blockage of Ca^2+^ currents due to Pb^2+^ interactions with ion selectivity filters.^14, 15^ Another example is the ligand-gated ionotropic N-methyl D-aspartate receptor (NMDAR),^16, 17^ where Pb^2+^ acts as an antagonist, partly through the interactions with the allosteric Zn^2+^ regulatory site^18^ in the extracellular domain of the receptor. An important class of Pb^2+^ targets are the intracellular Ca^2+^-sensor proteins, such as Synaptotagmin I (SytI),^19^ Calmodulin (CaM),^20, 21^ and protein kinase C (PKC).^22, 23^

While the proteins in question are quite distinct in their structure and function, one shared feature is the prevalence of oxygen donor ligands in their metal-ion coordination sites. The proposed NMDAR Pb^2+^-binding site comprises the oxygens of aspartate and glutamate carboxylate groups, along with additional nitrogen ligands provided by histidine residues.^24^ The selectivity filters of Ca^2+^ channels,^14, 25^ the EF hand motif of CaM,^21^ and the loop regions of the Ca^2+^-dependent phospholipid-binding conserved homology 2 (C2) domains of SytI^19, 26^ and PKC^23^ have all-oxygen metal-ion coordination sites capable of interactions with Pb^2+^. The analysis of Pb^2+^-bound protein structures in the PDB revealed that about 79% of the Pb^2+^-coordinating ligands are oxygen atoms that belong to the sidechain carboxylate and backbone carbonyl moieties of proteins, in addition to surrounding water molecules.^27^ The objective of this work was to determine what makes Pb^2+^ an effective competitor for oxygen-rich coordination sites in proteins, using SytI as a paradigm.

SytI is an integral membrane protein that serves as a Ca^2+^-dependent trigger of synchronous neurotransmitter release.^28^ The N-terminal segment of SytI is a transmembrane helical domain that anchors the protein to synaptic vesicle (**Fig. 1a**). The cytosolic C-terminal region comprises two Ca^2+^-sensing C2 domains, C2A and C2B. These domains have tri-partite (C2A) and bipartite (C2B) Ca^2+^ binding motifs that are believed to be targeted by Pb^2+^ with unknown stoichiometry.^19, 26^ The intrinsic Ca^2+^ affinities are pH-dependent and weak, ranging from 50 μM >10 mM.^29–31^ Ca^2+^ binding generates a localized electropositive potential in the apical C2 loop region and thereby enables SytI to interact with presynaptic membranes and SNARE proteins (**Fig. 1a**).^29, 32–36^ The outcome is the exocytic membrane fusion with the concomitant release of neurotransmitters into the synaptic cleft.

In this work, we demonstrate that SytI has two high-affinity Pb^2+^ binding sites, one per C2 domain. These high-affinity interactions, combined with fast binding and slow dissociation, impart thermodynamic and kinetic advantage on Pb^2+^ compared to Ca^2+^. Moreover, a single Pb^2+^ ion binding to either C2 domain has a profound inhibitory effect on subsequent Ca^2+^ binding, despite the existing coordination vacancies. Together, the inhibition of Ca^2+^ binding and previously known ability of Pb^2+^ to trigger membrane association of C2 domains^19, 23, 37^ provide a potential mechanism to explain the effect of Pb^2+^ on neurotransmitter release.

## METHODS

### Materials

The working solutions of metal ions were prepared in HPLC grade water or decalcified buffers using the following salts: Pb(II) acetate tri-hydrate (Sigma-Aldrich), standardized 1 M solution of Ca(II) chloride (Fluka Analytical), and Tb(III) chloride hexahydrate (Acros Organics). Prior to use, all buffers were treated with the ion-chelating resin Chelex 100 (Sigma-Aldrich) to remove trace divalent metals. Lipid components used in the phospholipid vesicle preparations: 1-palmitoyl-2-oleoyl-sn-glycero-3-phosphocholine (POPC) and 1-palmitoyl-2-oleoyl-sn-glycero-3-phospho-L-serine (POPS) were obtained from Avanti Polar Lipids Inc. (Alabaster, AL). The quartz cuvettes used for the Tb^3+^ luminescence experiments were coated with Sigmacote^®^ to avoid the protein adhesion to the walls. The cDNA of murine Syt1 was purchased from Open Biosystems (GE Life Sciences). All protein constructs were expressed and purified as described in the SI.

### Crystallization, structure determination and refinement

The samples used for crystallization of SytI domains with Pb^2+^ contained: (i) 17 mg/mL C2A with 7 mM Pb(II) acetate, and (ii) 22 mg/mL C2B with 1.1 mM Pb(II) acetate in 20 mM MES buffer at pH 6.0. Screening for crystallization was carried out in automated manner by using the sitting drop vapor-diffusion method with an Art Robbins Instruments Phoenix system in the X-ray Crystallography Core Laboratory at UTHSCSA. Crystals for Pb^2+^-bound C2A were obtained from Qiagen Classics II Suite condition #74 (0.2 M lithium sulfate, 0.1 M bis-tris pH 5.5, 25% polyethylene glycol 3350) at 4 °C. Although C2B was loaded with Pb^2+^ prior to crystallization, it was difficult to produce a Pb^2+^-loaded C2B crystal as the metal was typically lost resulting in apo-C2B crystals. Crystals for Pb^2+^-bound C2B were ultimately obtained from Microlytic MCSG-2 Suite condition #33 (0.2 M sodium fluoride, 20% polyethylene glycol 3350) at 22 °C. The crystals exhibited low occupancy Pb^2+^-binding during refinement of the structure coordinates, so an additional crystal was soaked overnight in mother liquor containing 5 mM lead acetate. This technique was applied to promote complete Pb^2+^-binding since the unsoaked crystal structure showed ambiguity in some of the electron density containing the binding site. The details of structure determination and refinement are given in the SI, along with the data collection and refinement statistics (**Table S1**). The refined coordinates of the Pb^2+^ complexes of C2A and C2B were deposited in the Protein Data Bank under accession codes 5vfe and 5vfg (5vff for partial Pb^2+^ occupancy), respectively. The analysis of metal-oxygen distances and the calculation of backbone r.m.s.d. from the previously published SytI structures (**Tables S2-S5**) was conducted using UCSF Chimera.^38^

### Isothermal Titration Calorimetry (ITC)

For ITC experiments, the C2A and C2B domains of SytI were extensively dialyzed against the large excess of decalcified ITC buffer (20 mM MES at pH 6.0 with 150 mM KCl). The filtered and degassed dialysis buffer was then used to prepare 50 μM C2A/C2B and 0.5 mM Pb^2+^ working solutions. The measurements for the heat of binding were carried out in MicroCal iTC200 (Malvern Panalytical) instrument with 14 successive additions of Pb^2+^ stock solution (0.5 μl for the first injection and 3 μl for all subsequent injections) into the protein. The acquisition and analysis of the triplicates was done using Origin software; the data were fit into a single set-of-sites binding model.

### Nuclear Magnetic Resonance (NMR) spectroscopy

#### NMR-detected Pb^2+^ binding to C2 domains

NMR-detected Pb^2+^ binding to [U-^15^N] enriched SytI C2 domains was monitored by acquiring series of ^1^H-^15^N HSQC spectra at 25 °C on Bruker AVANCE III spectrometers operating at ^1^H Larmor frequencies of 500 MHz (C2B) and 600 MHz (C2A). Protein concentration of 100 μM in decalcified 20 mM MES buffer (pH 6.0), 0.02% NaN_3_, and 8% D_2_O was used for all binding experiments. The Pb^2+^ concentrations were: 0, 0.025, 0.05, 0.075, 0.1, 0.125, 0.15, 0.2, 0.3, 0.4, 0.5, 0.75, 1.0, 1.3, 1.6, 2.0, and 2.5 mM for C2A; and 0, 0.0125, 0.025, 0.05, 0.075, 0.1, 0.125, 0.15, 0.2, 0.25, 0.3, 0.4, 0.5, 0.6, 0.8, 1.5, and 3.0 mM for C2B. The spectra were processed using NMRPipe^39^ and analyzed using Sparky.^40^ The chemical shift perturbation (CSP) due to M^2+^ binding, Δ, was calculated using the following equation:

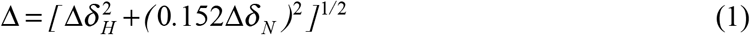

where Δδ_H_ and Δδ_N_ are residue-specific ^1^H and ^15^N chemical shift differences between the apo and Pb^2+^ bound states of the proteins. Pb^2+^ binding curves for the second site of the respective domains were constructed by plotting Δ as a function of corrected Pb^2+^ concentration to take into account the partial occupancy of the first metal binding sites. The binding curves were globally fitted (12 C2A, and 9 C2B residues) with a single-site binding model:

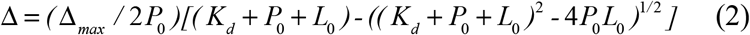

where Δ is the CSP value between the apo and Pb^2+^-bound state; Δ_*max*_ is the maximum CSP value reached upon Site 2 saturation; and *P*_*0*_ and *L*_*0*_ are the total protein and Pb^2+^ concentrations, the latter corrected for Pb^2+^ populating Site 1.

#### ZZ exchange NMR spectroscopy

The kinetic parameters of Pb^2+^ binding to the high-affinity sites of [U-^15^N] enriched SytI C2 domains were obtained by acquiring a series of ZZ exchange experiments^41^ on the cryoprobe-equipped Bruker AVANCE III spectrometers operating at ^1^H Larmor frequencies of 800 MHz (C2A) and 600 MHz (C2B). The data were collected at 4 different temperatures: 10, 15, 20, and 25 °C. The temperatures were calibrated using deuterated methanol. The protein samples (350 μM) were prepared in decalcified 20 mM MES buffer (pH 6.0), 150 mM KCl, 0.02% NaN_3_, and 8% D_2_O. Pb^2+^ was added to a concentration of 175 μM to generate approximately equal populations of the apo-and Pb^2+^-bound proteins. The samples were equilibrated overnight. The exchange between the two states, apo and Pb^2+^-bound, resulted in the transfer of longitudinal ^15^N magnetization during the variable mixing time period, manifested as the build-up of the cross-peak intensities and decay of the auto-peak intensities. The respective build-up and decay of the cross-peak and auto-peak intensities for the well-resolved residues was quantified as a function of effective mixing times: 12.53, 17.53, 22.53, 27.53, 32.53, 37.53, and 42.53 ms for C2A and an additional point of 52.53 ms for C2B (**Fig. S3**). Effective mixing times, *t*_*mix*_, were calculated as the duration of the mixing period plus 12.53 ms, which the time that ^15^N magnetization was longitudinal during the other elements of the pulse sequence. The spectra were processed using NMRPipe^39^ and analyzed using Sparky.^40^ The analysis of the ZZ exchange data was conducted as described in the SI, following the formalism of Miloushev et al.^42^

### Detection of mixed Pb^2+^/Ca^2+^ C2 species

The formation of C2·Pb1·(Ca)n complexes (n=1,2 for C2A and n=1 for C2B) was monitored at 25 °C on Bruker AVANCE III spectrometers operating at ^1^H Larmor frequencies of 800 MHz (C2A) and 500 MHz (C2B). The C2 domains were first equilibrated with concentrations of Pb^2+^ sufficient to selectively populate first metal binding sites in a buffer solution composed of decalcified 20 mM MES (pH 6.0), 150 mM KCl, 0.02% NaN_3_, and 8% D_2_O. The aliquots of Ca^2+^ solution prepared in the NMR buffer were added to the samples to achieve concentrations ranging from 100 μM to 40 mM. The spectral changes were monitored using ^15^N-^1^H HSQC spectra.

### *Tb*^*3+*^ Luminescence And Vesicle Co-Sedimentation Experiments Are Described In The Si

## RESULTS AND DISCUSSION

### Pb^2+^ targets Ca^2+^ Site 1 in both C2 domains of SytI

Previous studies of SytI suggested that Pb^2+^ binding site resides on the C2A domain.^19^ It was unclear to us from the structural viewpoint why C2A would be a preferred interaction site over C2B. To understand the structural basis of C2-Pb^2+^ interactions in SytI, we determined two high-resolution crystal structures of Pb^2+^-complexed C2A and C2B domains. Both structures revealed the presence of a single Pb^2+^ ion that was refined at high occupancy: C2A·Pb1 (1.4 Å, 5vfe) and C2B·Pb1 (1.8 Å, 5vfg) (**Table S1** and **Fig. 1b**). We found that the position of bound Pb^2+^ ion (Pb1) coincides with that of the first Ca^2+^ ion (subsequently referred to as Ca1, see **Fig. 1e,f**), as defined in previous structural studies of Ca^2+^-complexed C2 domains. Comparison of Pb^2+^-bound and apo structures showed that the most conformational changes due to Pb^2+^-binding are in the apical loop regions, specifically loop 1 in C2A and loop 3 in C2B (**Fig. 1b**). Close inspection of the Pb^2+^ coordination sites revealed that these differences are due to the rotation of the coordinating aspartic acid sidechains towards the metal ion: Asp178 in C2A, and Asp365 and Asp303 in C2B (**Figs. 1c,d**). The conformational changes in the other regions of C2 domains are minor, as evidenced by the low r.m.s.d. values obtained from the comparative analysis of existing C2A and C2B structures (**Tables S2-S3**).

Pb^2+^ ions bound to the C2A and C2B domains have a coordination number (CN) of 8. All ligands are oxygen atoms donated by the aspartic acid sidechains, one backbone carbonyl group, and one (or two in case of C2A) water molecules (**Figs. 1c,d**). The ligands are distributed uniformly in the coordination sphere, indicating that the 6s^2^ lone pair of Pb^2+^ is stereo-chemically inactive. The distribution of Pb-oxygen bond distances is narrow, ranging from 2.4 to 2.8 Å (**Tables S4-S5**). In coordination chemistry of Pb^2+^, uniform distribution of ligands and narrow range of Pb-ligand distances define a holodirected coordination geometry, which is favored in Pb^2+^ sites with high CN values and bulky ligands.^43^

**Figure 1.**
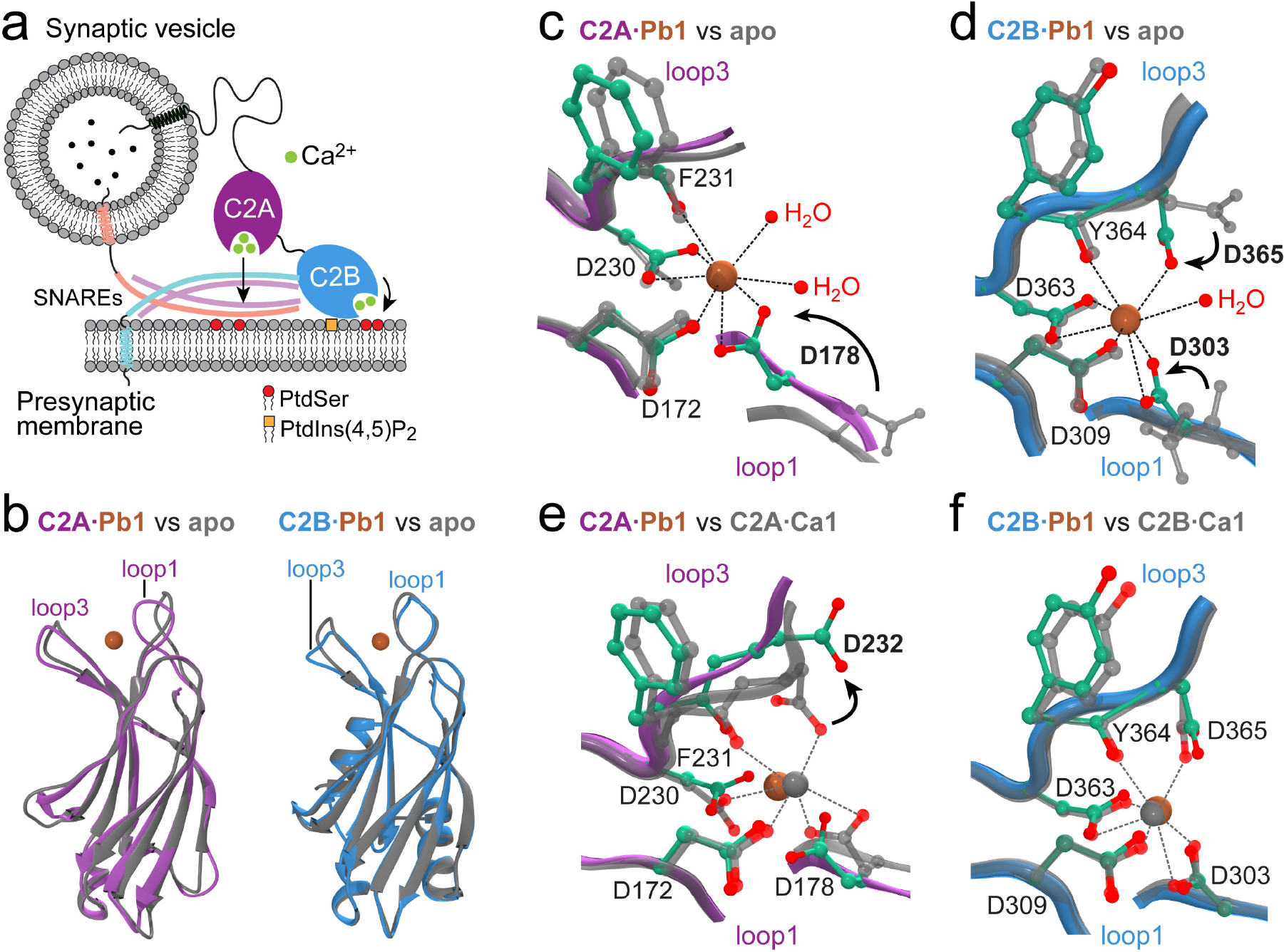
Structural analysis of Pb^2+^-SytI interactions. (**a**) SytI is a Ca^2+^-dependent trigger of exocytic membrane fusion. Ca^2+^ binding to C2A and C2B domains drives their interaction with anionic phospholipids of the presynaptic membrane, PtdSer and PtdIns(4,5)P_2_. (**b**) Crystal structures of Pb^2+^-complexed C2 domains reveal a single bound Pb^2+^ ion (sienna). Backbone superposition of Pb^2+^-complexed (C2A, purple and C2B, blue) and apo C2 domains (gray) illustrates the extent of conformational changes in the backbone of loop regions. The PDB identifiers are: 5vfe (C2A·Pb1), 4wee (apo C2A), 5vfg (C2B·Pb1), and 5ccj (apo C2B). (**c,d**) Octa-coordinated geometry of C2-bound Pb^2+^. The sidechain carbons and coordinating oxygens in the Pb^2+^-complexed structures are shown in green and red, respectively. All ligands are oxygen atoms donated by protein and water. Pb^2+^ binding is accompanied by the conformational rearrangement of several coordinating residues that are shown in boldface. (**e,f**) Comparison of metal-ion coordination sites in Pb^2+^- and Ca^2+^-complexed C2A (1byn/NMR) and C2B (1tjx) domains. Pb^2+^ and Ca^2+^ are represented with sienna and gray spheres, respectively. The coordination geometry of Ca^2+^ is shown with dashed lines. Only Ca1 and protein ligands are shown for clarity.

One notable difference between the Pb^2+^- and Ca^2+^-complexed C2A structures is a lack of coordination bond between Pb1 and the sidechain oxygens of Asp232, which points away from the metal ion binding site (**Fig. 1e**); in contrast, the Ca^2+^-Asp232 Oδ1 coordination bond is present in the NMR structure of C2A (1byn).^44^ In C2B, the coordination geometry of Ca1 and Pb1 is identical, which is also reflected in the similarity of the backbone conformation of the loop regions.

### Each C2 domain has one tight and one weak Pb^2+^ site

C2A and C2B have tri- and bi-partite Ca^2+^-binding motifs, respectively. To determine how many sites Pb^2+^ populates in solution, we conducted NMR-detected binding experiments of Pb^2+^ to the C2 domains. The chemical shift changes of the N-H_N_ backbone groups proximal to the M^2+^ coordination centers revealed two distinct Pb^2+^ binding events in C2A (**Fig. 2a**) and C2B (**Fig. 3a**). The first Pb^2+^ binding event, which is “slow” on the NMR chemical shift timescale for the majority of residues and saturates at approximately stoichiometric concentrations of Pb^2+^, gives rise to two sets of cross-peaks that correspond to the apo C2 and the C2·Pb1 complex. The second binding event is “fast”, manifesting itself in the smooth cross-peak trajectories as a function of increasing total Pb^2+^ concentration. These data indicate the presence of two Pb^2+^ sites with distinct kinetics and thermodynamics of binding. We did not observe an appreciable population of Site 3 by Pb^2+^ in the C2A domain.

To determine the influence of Pb^2+^ binding on the C2 loop regions, we constructed the chemical shift perturbation (CSP) plots for the high and low Pb^2+^ concentration regimes (**Fig. 2b** and **Fig. S1b**). The low concentration regime mostly reflects protein response to binding event 1, while the high concentration regime reflects the response to binding event 2. The CSP plot of C2B shows that all three loop regions are affected by interactions with Pb^2+^, with the most changes caused by the first binding event (**Fig. S1b**). In C2A, the second binding event influences the residues of loop 3 more than those of loop 1 (**Fig. 2b**). Using this information in conjunction with the Ca^2+^-bound C2A structure, we can then assign the low-affinity Pb^2+^ site to Site 2 and high-affinity Pb^2+^ site to Site 1 in C2A, which is populated in the crystal structure of the Pb^2+^ complex.

**Figure 2.**
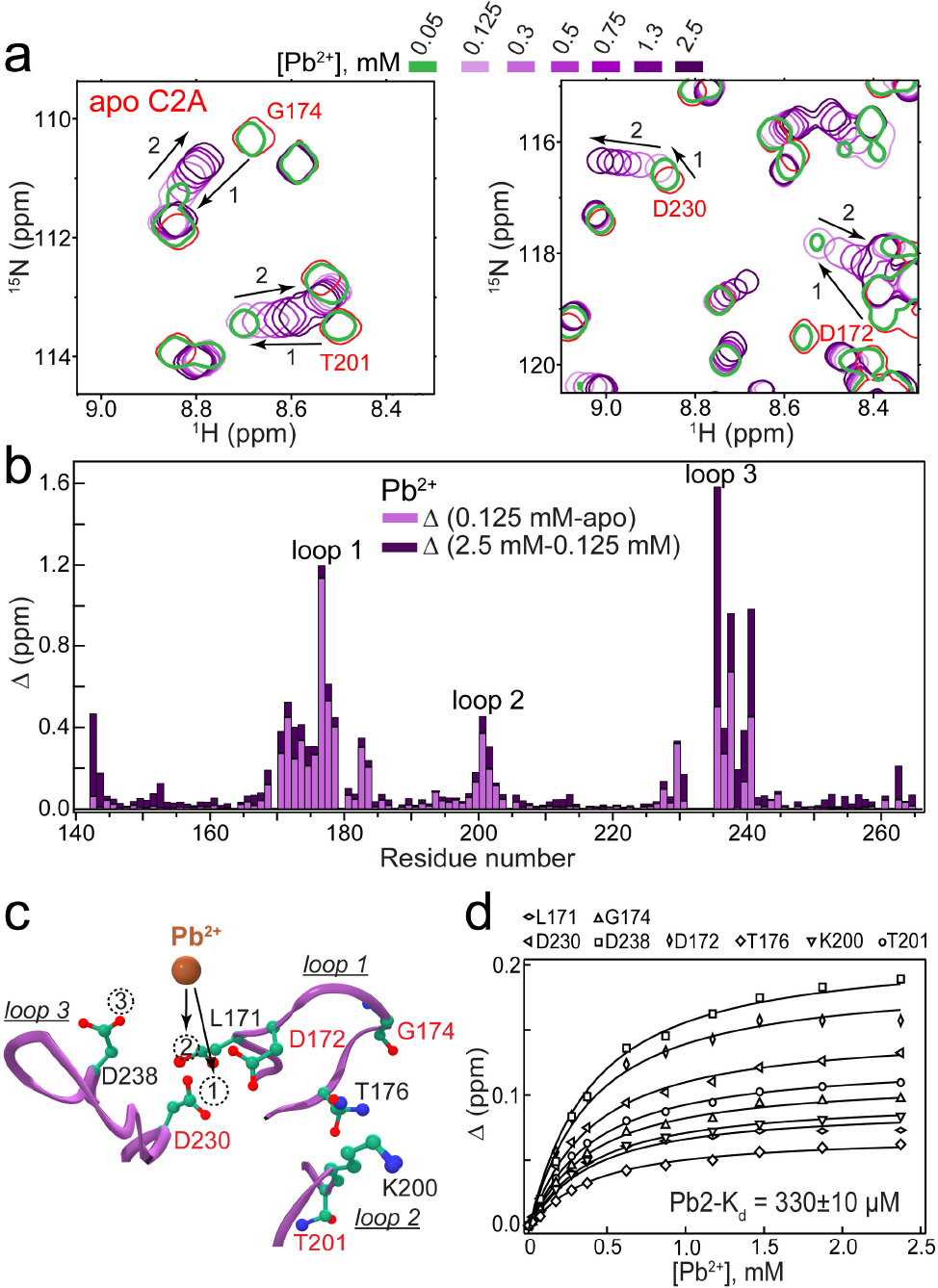
C2A domain binds two Pb^2+^ ions in solution. (**a**) Expansions of the C2A ^15^N-^1^H HSQC region for Pb^2+^ concentrations ranging from 0 to 2.5 mM. Peak displacements due to binding Pb1 (1, slow exchange) and Pb2 (2, fast exchange) are shown with arrows. (**b**) C2A chemical shift perturbation plot constructed for the low- (< 0.125 mM) and high- (> 0.125 mM) concentration regimes of Pb^2+^. (**c**) Loop region of the C2A showing the location of Ca^2+^ sites 1-3, along with residues whose ^15^N-^1^H cross-peaks are labelled in (**a**) and/or used in (**d**). (**d**) NMR-detected binding curves constructed for the low-affinity Pb^2+^ Site 2. Solid lines represent the global fit that produced the K_d_ of 330 ± 10 μM.

We used the fast-exchange NMR data to construct the binding curves and obtain Pb^2+^ affinities to Site 2 of the C2 domains (see **Fig. 2c** and **Fig. 3b** for the Site 2 location). Global fitting of the binding curves produced *K*_*d*_ values of 330 ± 10 μM (C2A, **Fig. 2d**) and 220 ± 5 μM (C2B, **Fig. S1c**). The Ca^2+^ affinities for the same sites under identical buffer conditions are 1.6 mM (C2A) and 0.7-0.8 mM (C2B).^31^ This means that the affinity of Pb^2+^ to Site 2 exceeds that of Ca^2+^ by 5- and 3-fold in the C2A and C2B domains, respectively.

**Figure 3.**
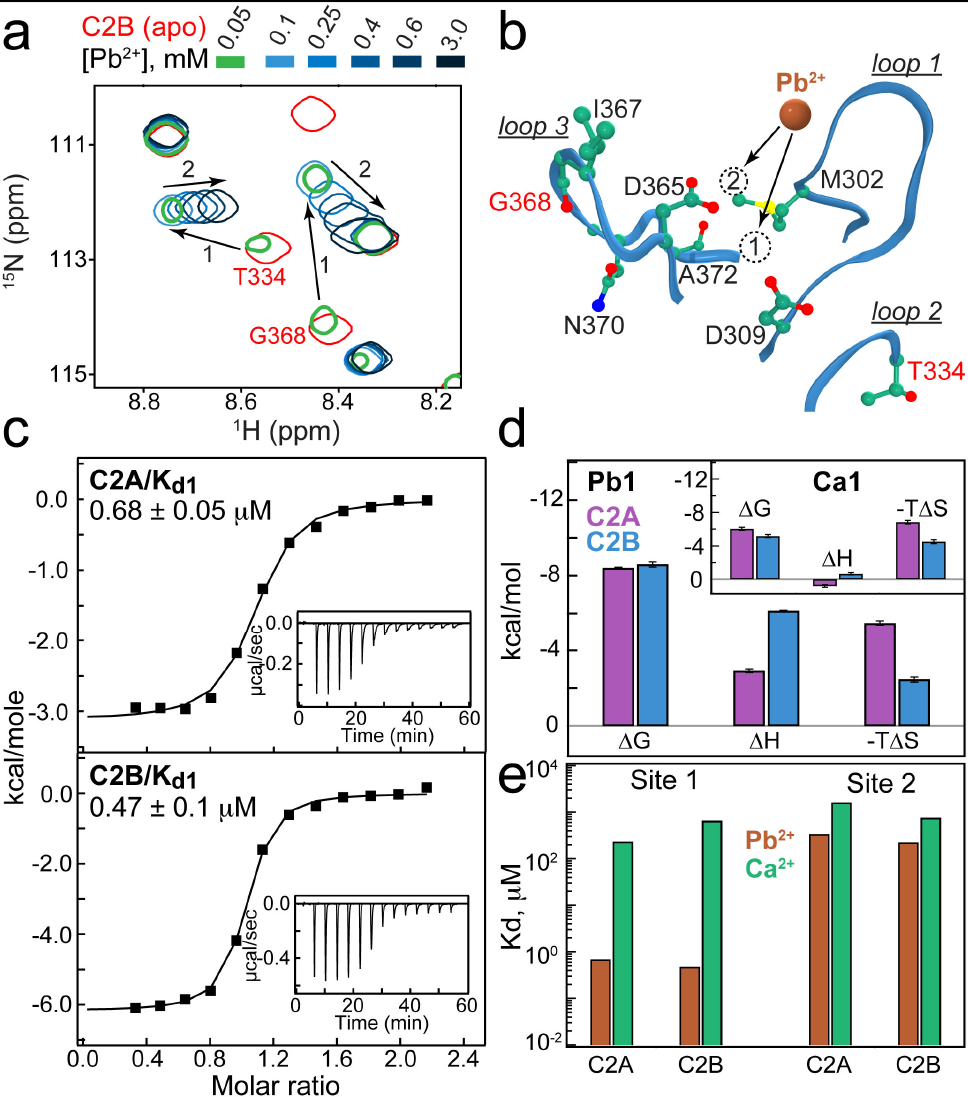
C2 domains bind two Pb^2+^ ions in solution, with differential affinities. (**a**)Expansions of C2B ^15^N-^1^H HSQC region for Pb^2+^ concentrations ranging from 0 to 3 mM. Peak displacements due to binding Pb1 (1, slow exchange) and Pb2 (2, fast exchange) are shown with arrows. (**b**) Loop region of C2B showing the location of Ca^2+^ sites 1 and 2 that are populated by Pb^2+^ in solution. (**c**) ITC profiles for Pb1 association with C2A (top) and C2B (bottom), and the respective dissociation constants. (**d**) Thermodynamic parameters of Pb1 binding to C2 domains. The inset shows Ca1 data reported in the previous study.^45^ (**e**) Comparison of Pb^2+^ and Ca^2+^ affinities to sites 1 and 2 measured under identical conditions.

The slow exchange regime displayed by Pb^2+^ binding to Site 1 is generally unsuitable for the determination of binding affinities. We therefore conducted ITC experiments to obtain the dissociation constants (*K*_*d*_) and thermodynamic parameters of Pb^2+^ binding to Site 1. The *K*_*d*_ values of Pb1 are in the sub-micromolar range for both domains: 0.68 ± 0.05 μM (C2A) and 0.47 ± 0.1 μM (C2B), respectively (**Fig. 3c**). This represents 340-fold (C2A) and 1,400-fold (C2B) enhancement of Pb^2+^ affinities compared to those of Ca^2+^ under identical buffer conditions.^31^ The underlying thermodynamic basis of this enhancement is evident from the comparison of the enthalpic and entropic contributions to Δ*G* (**Fig. 3d**). Pb1 binding to C2 domains is significantly exothermic. Combined with the favorable entropic contribution, this leads to large negative Δ*G* values. In contrast, the enthalpic contribution to Ca1 binding is small (inset of **Fig. 3d**), with Δ*G* being dominated by the entropy term. Therefore, it is mostly the differences in binding enthalpies that are responsible for the differential affinities of Pb1 and Ca1. The positive entropy change for both metal ions indicates that the gain due to metal de-solvation^46^ is sufficient to compensate for the loss of conformational flexibility of metal ion-coordinating ligands.

A comparative summary of the binding data illustrates the two main conclusions of our experiments (**Fig. 3e**). First, Pb^2+^ populates Sites 1 and 2 in solution, with Pb1 affinity exceeding that of Pb2 by ~500-fold; this property enabled us to selectively probe Pb^2+^ binding events using ITC and NMR. Second, Pb^2+^ affinities are higher than those of Ca^2+^ for both C2 domains. The enhancement of Pb^2+^ affinities compared to those of Ca^2+^ is significantly more pronounced for Site 1, which is the only site populated in the crystal structures of C2A and C2B.

Our conclusions regarding differential affinities of Pb^2+^ and Ca^2+^ to the C2 domains of SytI are further supported by the results of Tb^3+^ displacement experiments (**Fig. 4**). Tb^3+^ binds to the cytoplasmic region of SytI that contains both C2A and C2B domains (C2AB) with an apparent affinity of 2.5 μM (data not shown). When bound to C2AB, Tb^3+^ shows a strong luminescence signal due to FRET from the tryptophan sidechains. We prepared a fully Tb^3+^-saturated C2AB and monitored the intensity changes of the strongest luminescence peak at 545 nm upon addition of Ca^2+^ and Pb^2+^ (**Fig. 4a,b**). It takes ~100-fold more Ca^2+^ than Pb^2+^ to achieve ~50% Tb^3+^ displacement from C2AB (**Fig. 4c**), clearly indicating the thermodynamic preference for Pb^2+^ over Ca^2+^.

**Figure 4.**
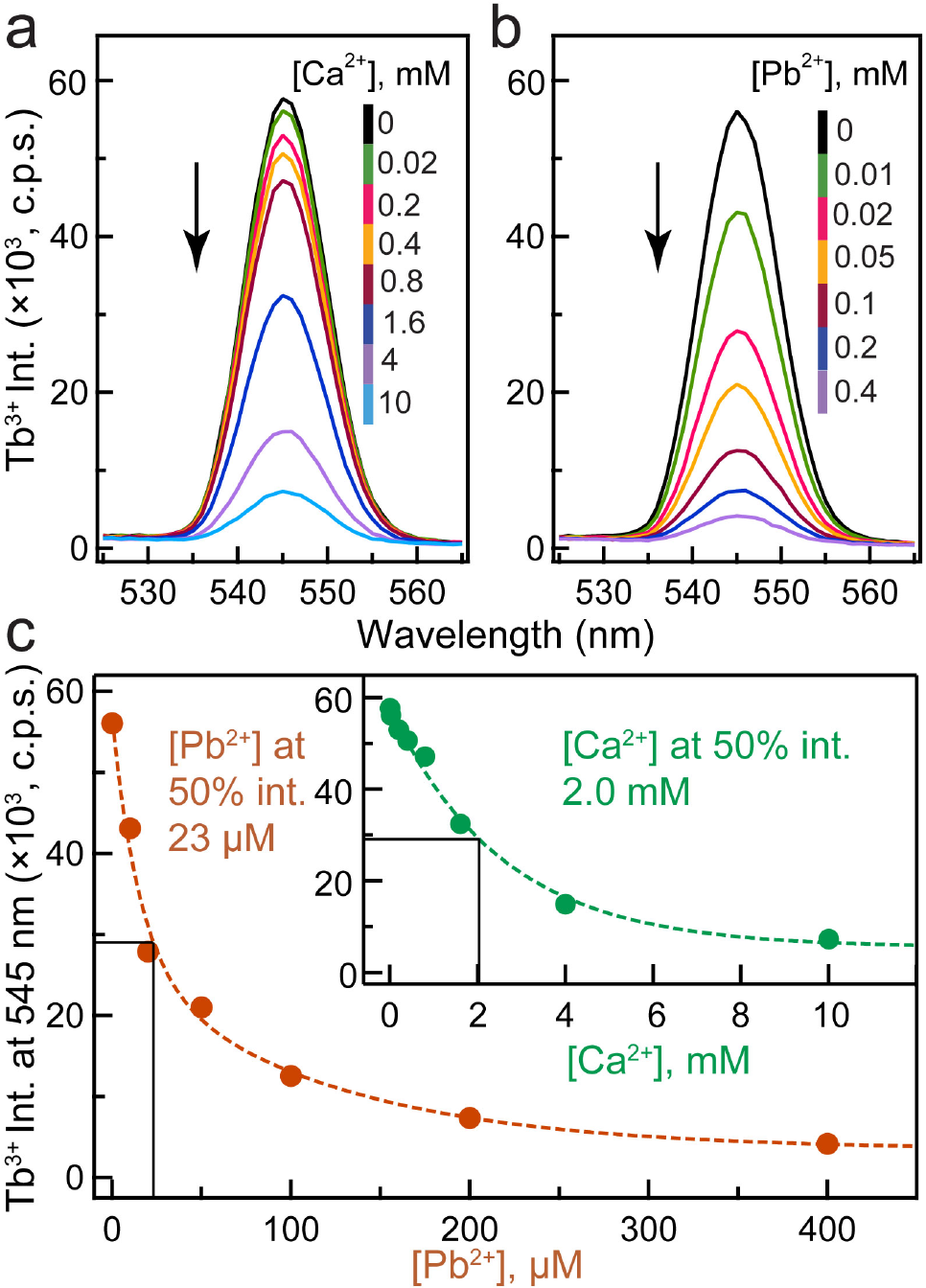
Pb^2+^ is more potent than Ca^2+^ in displacing Tb^3+^ from the SytI C2AB region. The most intense Tb^3+^ emission peak at 545 nm (^5^D_4_ to ^7^F_5_ transition) is shown as a function of increasing Ca^2+^ (**a**) and Pb^2+^ (**b**) concentrations. The decrease in luminescence is indicative of the displacement of Tb^3+^ from the protein by Ca^2+^ and Pb^2+^. (**c**) Intensity decrease at 545 nm plotted as a function of M^2+^ (M=Ca, Pb).

It is well established that the affinities of divalent metal ions to C2 domains significantly increase in the vicinity of anionic lipids, in what Falke coined as a “target-activated messenger affinity” (TAMA) mechanism.^47^ The implication is that intrinsic metal-ion affinities that we measure in solution are 2-3 orders of magnitude lower than those at the membrane surface. This mechanism provides an explanation of why C2 domains, being intrinsically weak Ca^2+^-binding modules in solution, are able to respond to low micromolar Ca^2+^ concentrations during the cell-signaling event. In the framework of the TAMA mechanism, the affinity of Pb^2+^ to Site 1 would be comparable to the bioavailable concentration of Pb^2+^, which ranges from picomolar to nanomolar.^48^ This would make Pb^2+^ binding feasible under physiological conditions. The role of Ca^2+^ binding to Site 1 in C2 domains is the initial weak recruitment of the protein complex to the membranes, with the subsequent population of remaining Ca^2+^ site(s) to ensure high-affinity interaction.^47^ Consistent with these findings, we observed very weak interactions between the C2·Pb1 complexes and phosphatidylserine-containing LUVs (data not shown), but almost full membrane association of C2 under saturating Pb^2+^ conditions (**Fig. S2**). Based on the above considerations, we conclude that: (i) Pb^2+^ is isofunctional to Ca^2+^ in its ability to support the C2-membrane interactions; and (ii) interactions of Ca^2+^ and Pb^2+^ with Site 1 of C2 domains will primarily determine the competitive behavior of these metal ions. We next sought to explore the properties of the C2·Pb1 complexes that – in addition to relative Ca1 and Pb1 affinities (**Fig. 3e**) – are most relevant for Pb^2+^/Ca^2+^ competition: the kinetics of Pb^2+^ binding to Site 1 and the formation of mixed metal ion C2 species.

### Fast binding and slow dissociation of Pb^2+^ from Site 1

To obtain the kinetic information on Pb^2+^ binding to Site 1 of the C2A and C2B domains, we used the ZZ-exchange solution NMR spectroscopy. The method relies on the exchange of longitudinal magnetization between two C2 domain species: apo and single Pb^2+^-bound, C2·Pb1 (**Fig. 5a,b**). The inter-conversion between apo C2 and C2·Pb1 gives rise to cross-peaks (inset of **Fig. 5c,d**). The time-dependence of the auto- and cross-peak intensities, expressed through the composite ratio Ξ,^42^ contains information on the on- and off-rate constants for Pb^2+^ binding to Site 1 (**Figs. S3-4** and **Fig. 5c,d**). The NMR data analysis produced the on-rate constants: (6.8 ±0.4) × 10^7^ M^−1^s^−1^ (C2A) and (6.2 ± 0.3) × 10^7^ M^−1^s^−1^ (C2B) that are close to the diffusion limit of 6 × 10^8^ M^−1^s^−1^ at 25 °C.^49^ Moreover, the Pb^2+^ *k*_*on*_ values are comparable to the *k*_*on*_ value previously reported for Ca1 binding to the C2A domain, (3.9 ± 0.8) × 10^7^ M^−1^s^−1^.^50^ In contrast to *k*_*on*_, the Pb^2+^ *k*_o*ff*_ values of 45.5 (C2A) and 29 s^−1^ (C2B) are approximately two orders of magnitude smaller than the 2000-9500 s^−1^ range previously reported for Ca^2+^.^50–52^ Therefore, the differential affinities of Ca^2+^ and Pb^2+^ to Site 1 are due to the differences in the off-rate constants.

The temperature-dependent kinetics data were further used to estimate the activation enthalpy Δ*H*^≠^ and activation entropy Δ*S*^≠^ for the forward and reverse reactions (**Fig. 5e**). Although the enthalpic barrier Δ*H*^≠^_f_ for the C2A-Pb^2+^ association (9.3 kcal/mol) is larger than that of C2B (6.6 kcal/mol), the accompanying differences in Δ*S*^≠^_f_ values produce essentially identical Δ*G*^≠^_f_ values for C2A and C2B, 6.8 kcal/mol at 25 °C. This is only 1.8 kcal/mol larger than the theoretically predicted energy cost of ~5 kcal/mol required to transport a small molecule at the diffusion limit.^49^ The negligible Δ*S*^≠^_f_ for the C2B-Pb^2+^ association indicates that the gain in solvent entropy due to de-solvation of Pb^2+^ and protein binding region is offset by a loss of conformational flexibility of C2B in the transition state. This is in contrast to C2A, where the positive value of Δ*S*^≠^_f_ suggests that conformational flexibility in the transition state is partially preserved. In the reverse (dissociation) direction, the activation parameters for the C2A and C2B are similar, with enthalpy and entropy terms contributing 80% and 20% to Δ*G*^≠^_r_, respectively. In aggregate, our data suggest that Pb^2+^ can act as both a thermodynamic and kinetic trap for the C2 domains and thereby effectively compete with Ca^2+^ for Site 1.

**Figure 5.**
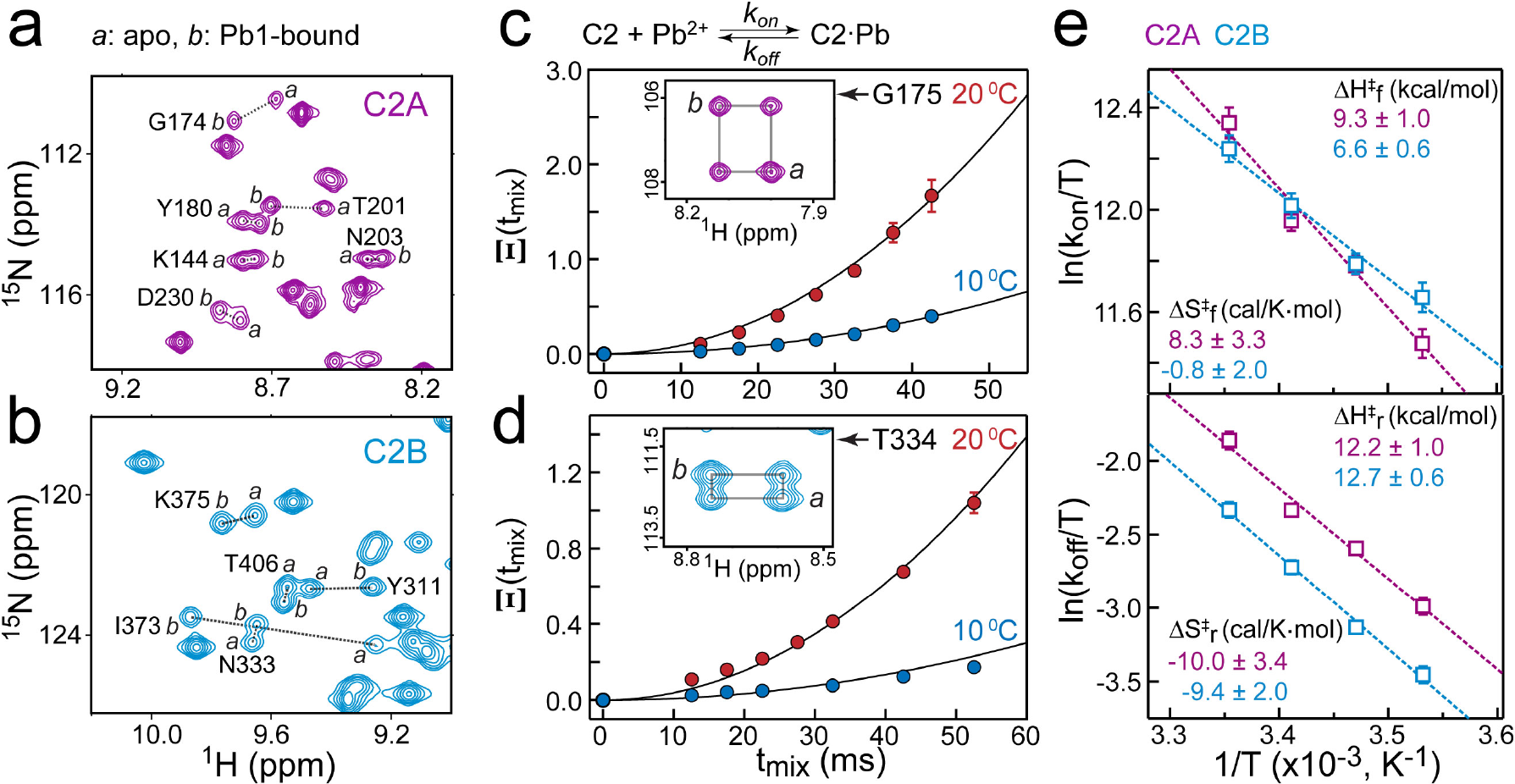
Kinetics of Pb^2+^ binding to Site 1 probed using ZZ exchange NMR spectroscopy. Slow exchange behavior between the apo and C2·Pb1 forms is illustrated using expansions of ^15^N-^1^H HSQC spectra of C2A (**a**) and C2B (**b**) domains. Representative ZZ exchange data, showing the time dependence of composite ratio Ξ at two different temperatures for Gly175 in C2A (**c**) and Thr334 in C2B (**d**). (**e**) Eyring plots for the temperature range (10-25 °C) accessible to ZZ exchange spectroscopy in this kinetic regime. The values of Δ*H*^≠^ and Δ*S*^≠^ were determined from the linear fit, which is shown with a dashed line.

### Pb^2+^ binding to Site 1 locks C2 domains in Ca^2+^-insensitive state

One potential mechanism through which Ca^2+^ can possibly rescue^53^ the Pb^2+^-induced protein behavior is through the formation of mixed metal ion species, with Pb^2+^ populating Site 1 and Ca^2+^ populating Site(s) 2 and 3, respectively (**Fig. 6a**). To test if this could be the case in SytI, we prepared C2·Pb1 complexes and evaluated their Ca^2+^-binding behavior in the Ca^2+^ concentration range from 0.1 to 40 mM, using solution NMR spectroscopy. The NMR samples were prepared such that the populations of C2A·Pb1 and C2B·Pb1 complexes were the dominant species (≥ 95 %), with negligible population of Site 2 by Pb^2+^. To our surprise, it took mM Ca^2+^ concentrations to detect noticeable shifts in the NMR spectra of C2·Pb1. Only at very high concentrations of Ca^2+^ (10-40 mM) did we observe a clear titratable Ca^2+^-dependent behavior of cross-peaks that belong to the loop residues (**Fig. 6b,c; Fig S5**).

**Figure 6.**
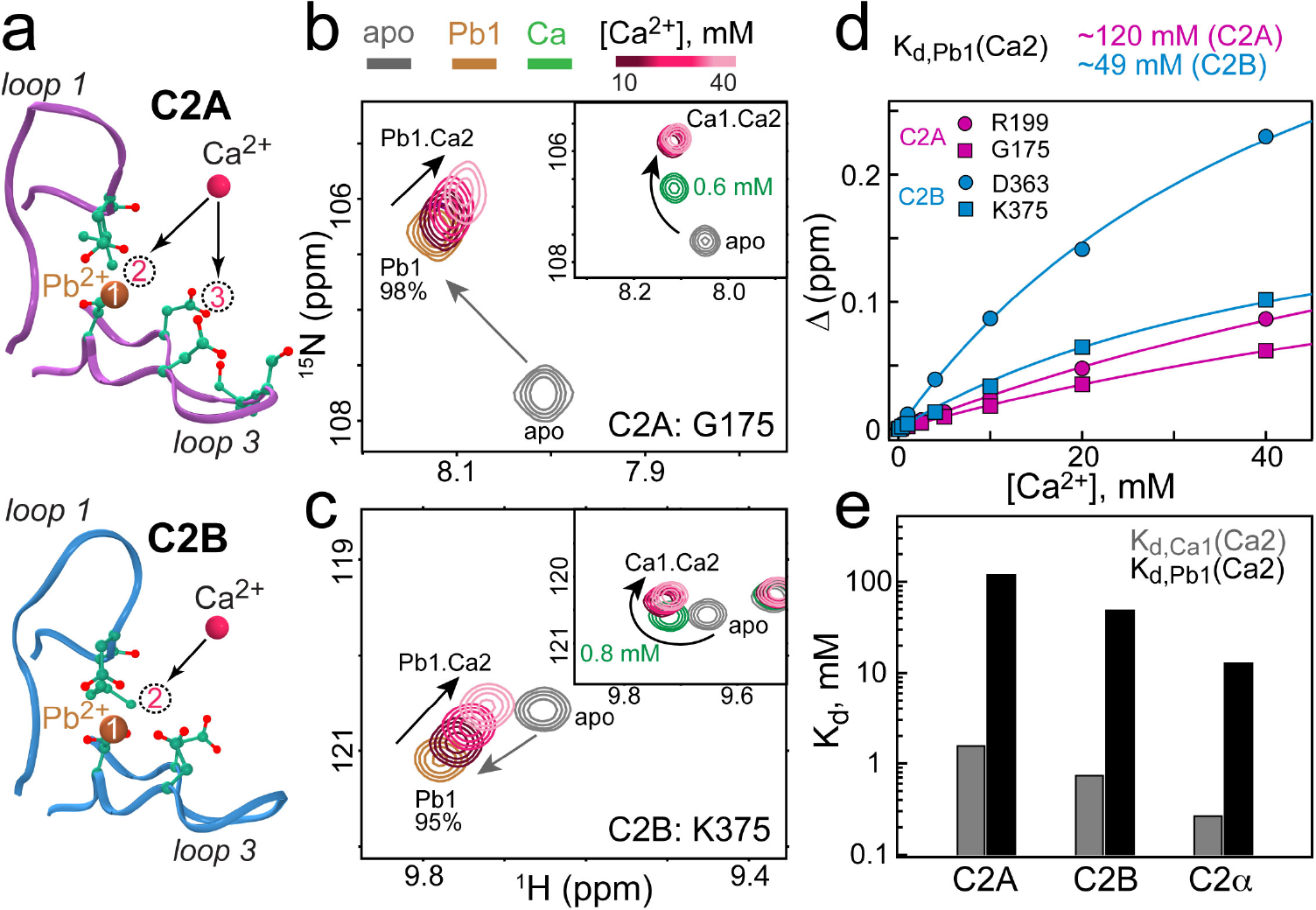
Pb^2+^ binding to Site 1 inhibits Ca^2+^ binding to the remaining coordination vacancies. (**a**) Location of metal ion coordination vacancies relative to Pb1 in the loop regions of C2A and C2B. (**b,c**) Expansions of the ^15^N-^1^H HSQC spectra showing Ca^2+^ binding behavior of Gly175 in C2A·Pb1 (b) and K375 in C2B·Pb1 (**c**). The C2 states are: apo (gray), C2·Pb1 (sienna), and C2·Ca1/Ca2 (green), the latter representing C2 domains at intermediate Ca^2+^ concentrations. The insets show cross-peak trajectories obtained in Ca^2+^-only binding experiments. While not even 40 mM Ca^2+^ is sufficient to saturate Site 2 of C2·Pb1 complexes, Site 2 saturates at <10 mM Ca^2+^ in the absence of pre-bound Pb^2+^. (**d**) Representative Ca^2+^ binding curves constructed for the C2·Pb1 complexes. Solid lines represent a global fit to the single-site binding equation that produced estimates of the dissociation constants. (**e**) Comparison of Ca^2+^ dissociation constants from Site 2 of C2A and C2B, when Site 1 is occupied by Ca^2+^ (gray) and Pb^2+^ (black).

This is in sharp contrast with the Ca^2+^-only binding data that showed full saturation of Sites 1 and 2 at ~10 mM Ca^2+^ (insets of **Fig. 6b,c**). The dissociation constants of Ca^2+^ from Site 2 of the C2A·Pb1 complexes, *K*_*d*,Pb1_(Ca2), obtained from the NMR data analysis are ~120 and ~49 mM for the C2A and C2B, respectively (**Fig. 6d**). Comparison of the *K*_*d*,Pb1_(Ca2) with previously reported *K*_*d*,Ca1_(Ca2) values^31^ indicates that population of Site 1 by Pb^2+^ reduces Ca^2+^ affinities to Site 2 by 60-70 fold. Moreover, the same pattern holds for the C2 domain from another protein, Protein Kinase Cα^23^ (**Fig. 6e**). Therefore, for three C2 domains that share about ~40% pairwise sequence identity we observed the same pattern: binding of Pb^2+^ to the high-affinity Site 1 on C2 domains reduces the Ca^2+^ affinity to the remaining vacant sites.

In view of the modest structural changes caused by Pb^2+^ binding to the C2 domain, the most likely explanation of this behavior lies in the electronic properties of Pb^2+^ influencing Ca^2+^ affinity through the “bridging” ligands. Inspection of the Ca^2+^-complexed structures of C2A and C2B shows that in each domain there are two oxygen atoms that bridge metal ions in Sites 1 and 2: Oδ1(Asp172) and Oδ1(Asp232) in C2A, and Oδ1(Asp303) and Oδ1(Asp365) in C2B (**Fig. S6a,b**).^44, 54^ Pb^2+^ is a stronger Lewis acid than Ca^2+^, which is manifested in its higher electronegativity.^55, 56^ A significant depletion of electron density of bridging ligands by Pb^2+^ at Site 1 would reduce their electron-donating abilities and result in the attenuation of Ca^2+^ affinities to Site 2. If we apply the same rationale to describe Ca^2+^ interactions with Site 3 of the C2A·Pb1 complex, then Ca^2+^ affinity should not be significantly affected because metal ions in Sites 1 and 3 do not share any oxygen ligands. Indeed, the *K*_*d*,Pb1_(Ca3) of 26 mM (**Fig. S6,c**) determined using our NMR data is not significantly different from the >10 mM estimate reported for the Ca^2+^-only system.^30^ Another important outcome of Ca^2+^-binding experiments is that we did not observe any evidence of Pb^2+^ displacement from Site 1 even at > 250-fold Ca^2+^ excess, further confirming our prediction that Pb^2+^ can effectively compete with Ca^2+^ for Site 1.

## CONCLUSIONS

We have characterized the structural, thermodynamic, and kinetic aspects of Pb^2+^ interactions with the C2 domains of SytI, a key regulator of the synaptic vesicle fusion and neurotransmitter release. We established that the Ca^2+^-binding Site 1 of the C2 domains is the most likely target of Pb^2+^ due to high affinity of the interactions. The slow dissociation kinetics of Pb^2+^ will increase the lifetime of the protein-Pb^2+^ complexes in the cell and thereby make Pb^2+^ a potent competitor of Ca^2+^. The most unexpected outcome of our study – the loss of Ca^2+^ sensitivity of the C2 domains when Pb^2+^ populates only a single high-affinity site – suggests possible mechanisms through which Pb^2+^, despite its low bioavailability, can disrupt the function of Ca^2+^-dependent proteins. For instance, the inhibition of Ca^2+^-dependent synchronous release of neurotransmitters^57–64^ observed upon Pb^2+^ exposure could be attributed to the failure of Pb^2+^-complexed SytI to sense elevated intracellular Ca^2+^ concentrations. In addition, the ability of Pb^2+^ to support the membrane interactions of SytI can explain the observed stimulatory effect of Pb^2+^ on Ca^2+^-independent spontaneous release. ^57–64^ Previously, Bouton et al^19^ demonstrated that Pb^2+^ is ~1000-fold more potent than Ca^2+^ in driving the membrane association of the cytoplasmic region containing both C2 domains of SytI. Combined with the results reported here, this offers an intriguing possibility that, in contrast to a full complement of Ca^2+^ ions, only one Pb^2+^ ion per C2 domain might be sufficient to drive the membrane interactions of SytI. Membrane-binding experiments on full-length SytI reconstituted into membrane-mimicking environment are required to address this question. Finally, our findings indicate that high-affinity interactions of Pb^2+^ with proteins are not limited to the thiol-rich coordination sites,^65–69^ but can also occur in the all-oxygen coordination environment provided by the C2 domains, the Ca^2+^-sensing membrane-binding modules found in many signaling proteins.

## Author contributions

T.I.I. and S.K. designed the study. T.I.I. directed the project. S.K. conducted the NMR spectroscopy, vesicle co-sedimentation, and luminescence experiments, along with the corresponding data analysis. B.H. and A.K.S. contributed to sample preparation and initial stages of the NMR and luminescence work. Samples for crystallization trials were prepared by B.H., A.K.S. and S.K. Structure determination by X-Ray crystallography was carried out by A.B.T. ITC data acquisition and processing were done by S.W.L. using protein samples prepared by S.K. T.I.I. and S.K. wrote the manuscript with input from all authors.

## Acknowledgement

This work was supported by the NSF CAREER Award CHE-1151435 to T.I.I. A.K.S. and S.K. were supported by the NIH Grant R01GM108998 (T.I.I.) and Welch Foundation Grant A-1784 (T.I.I.), respectively. The X-Ray Crystallography Core Laboratory is supported by the Office of the Vice President for Research and by the NIH/NCI Grant P30 CA054174 award to the Mays Cancer Center, the newly named center home to UT Health San Antonio MD Anderson Cancer Center. S.W.L acknowledges the support from Welch Foundation grant A-1742.

